# *Repetita iuvant:* repetition facilitates online planning of sequential movements

**DOI:** 10.1101/819938

**Authors:** Giacomo Ariani, Young Han Kwon, Jörn Diedrichsen

## Abstract

Beyond being essential for long-term motor-skill development, movement repetition has immediate benefits on performance, increasing speed and accuracy of a second execution. While repetition effects have been reported for single reaching movements, it has yet to be determined whether they also occur for movement sequences, and what aspects of sequence production are improved. We addressed these questions in two behavioral experiments using a discrete sequence production (DSP) task in which human volunteers had to perform short sequences of finger movements. In Experiment 1, we presented participants with randomly varying sequences and manipulated 1) whether the same sequence was repeated on successive trials, and 2) whether participants had to execute the sequence (Go), or not (No-Go). We establish that sequence repetition led to immediate improvements in speed without associated accuracy costs. The largest benefit was observed in the middle part of a sequence, suggesting that sequence repetition facilitated online planning. This claim was further supported by Experiment 2, in which we kept a set of sequences fixed throughout the experiment, thus allowing participants to develop sequence-specific learning: once the need for online planning decreased, the benefit of repetition disappeared. Finally, we found that repetition-related improvements only occurred for the trials that had been preceded by sequence production, suggesting that action selection and sequence pre-planning may not be sufficient to reap the benefits of repetition. Together, these results show that repetition can enhance representations at the level of movement sequences (rather than of individual movements) and facilitate online planning.

**New & Noteworthy:** Even for overlearned motor skills such as reaching, movement repetition improves performance. How brain processes associated with motor planning or execution benefit from repetition, however, remains unclear. Here we report the novel finding of repetition effects for sequential movements. Our results show that repetition benefits are tied to improved online planning of upcoming sequence elements. We also highlight how actual movement experience appears to be more beneficial than mental rehearsal for observing short-term repetition effects.

## Introduction

Repeated practice is an essential ingredient for motor learning. However, even the immediate repetition of the same stimulus, or response, often leads to better performance (i.e., “repetition effect”). Stimulus repetition enhances perceptual processing (Bentin and McCarthy 1994; Eichelman 1970) and improves stimulus-response (S-R) mapping (Bertelson 1961, 1963, 1965). Conversely, switching usually incurs a performance cost (Adams 1961; Eimer et al. 1995; Hyman 1953; Kleinsorge 1999; Smith 1968). Within the movement domain, repetition can provide short-term benefits to motor output (Vleugels et al. 2019), even for well-learned skills (Ajemian et al. 2010; Phatak et al. 2020). As a common example, athletes and musicians rehearse action sequences moments before a big match or performance. Previous research on reaching movements has shown that movements are biased towards the direction experienced in the recent history, and repeated movements can be executed with less variability (Chapman et al. 2010; Diedrichsen et al. 2010; Marinovic et al. 2017; Verstynen and Sabes 2011). Yet, the mechanisms by which movement repetition facilitates task performance remain elusive.

A recent study (Mawase et al. 2018) provided some insight into the possible origins of this effect. The authors argue that repetition accelerates movement pre-planning – the ability of the system to reach a well-prepared state, from which movements can be initiated and produced quickly and efficiently. The paper also presents some arguments that this effect was not caused by speeding up perceptual or action selection processes. However, many real-life motor skills are more complex than single, point-to-point reaches – they tend to involve the production of sequential movements. In this context, the general term planning can refer to either pre-planning, planning-related processes that occur prior to movement onset (during the preparation phase), or to online planning, planning-related processes that occur after movement onset (during the movement phase; Ariani and Diedrichsen, 2019). If repetition only improves the planning of individual movements, we should not find a repetition effect at the sequence level – that is, switching between two different orderings of the same movement elements should be as good as repeating the same ordering. Conversely, if repetition accelerates planning at the level of a sequence, we would expect to observe a repetition effect only when the ordering remains consistent.

Using a discrete sequence production (DSP) task, we have recently shown that faster performance for trained sequences relies on improvements in online planning – the ability to plan future elements in parallel with the execution of preceding sequence elements (Ariani and Diedrichsen 2019). Given the hypothesis that repetition improves movement planning (Mawase et al. 2018), we would therefore expect to observe a repetition benefit not only on reaction time, which depends on pre-planning and movement initiation, but also on sequence movement time, which depends on online planning and movement execution. To test these ideas, here we used a DSP task in which participants were explicitly cued to produce short sequences of finger movements (Exp. 1). On any given trial, the sequence could either be the same as in the previous trial (Repetition), or a different sequence (Switch), with equal probability (0.5). Participants were given enough time (2.5 s) to complete stimulus identification and action selection before the go signal. The use of such a delayed-movement paradigm ensured that repetition effects could not be caused by improved perceptual processes. Our findings indicated that sequence repetition improved both reaction times (the time between go signal and the first keypress) and sequence movement times (the time between first and the last keypress in the sequence).

Next, we asked the exploratory question of whether these benefits are caused by processes occurring before movement onset (stimulus identification and sequence pre-planning), or by processes occurring during sequence production (initiation, execution, and online planning). While our design encouraged participants to pre-plan each sequence during the preparation phase, we manipulated whether they had to perform the sequence, or not, with a Go/No-Go paradigm. This allowed us to compare Repetition and Switch trials, depending on the whether the previous (N-1) trial involved pre-planning alone (No-Go condition), or included also the initiation, execution, and online planning of the sequence (Go condition).

Finally, in a separate behavioral experiment on an independent sample of participants (Exp. 2), we examined how the repetition effect changed with the gradual development of sequence-specific learning by monitoring the effect over the course of training on a fixed set of sequences.

## Methods

### Participants

Forty-nine right-handed volunteers participated in Experiment 1 (33 F, 16 M; age 18–39, mean 22.73 years, SD 5.04). An independent sample of forty right-handed volunteers participated in Experiment 2 (24 F, 16 M; age 18–36, mean 22.28 years, SD 3.44). Handedness was assessed using the Edinburgh Handedness Inventory (Exp. 1: mean 86.33, SD = 15.67; Exp. 2: mean 78.59, SD = 16.74). While some participants had some musical training, none of them was a professional musician (musical experience, Exp. 1: mean 4.67 years, SD 6.01; Exp. 2: mean 3.14 years, SD 3.77). None of the participants had a history of neurological disorders. Experimental procedures were approved by the ethics committee at Western University (London, Ontario, Canada). All participants gave written informed consent and received monetary compensation for their participation. Four participants withdrew from Exp. 1 before study completion and were thus excluded from data analysis (final N = 45). In Exp. 2, one participant failed to follow the instructions. The session was terminated before completion and the data excluded from successive analysis (final N = 39).

### Apparatus

Sequences of finger presses were executed on a custom-made keyboard device comprised of five keys corresponding to each finger of the right hand. The isometric force exerted by each finger was continuously recorded by force transducers under each key (FSG-15N1A, Sensing and Control, Honeywell; dynamic range, 0-25 N) at a rate of 500 Hz. To account for sensor drifts, we recalibrated the zero-force baseline at the beginning of each block of trials. Each key was independently deemed to be *pressed* when the force exceeded a threshold of 1 N and *released* as soon as the force returned below 1 N. (Fig. 1C; P = press, R = release).

**Figure 1.**
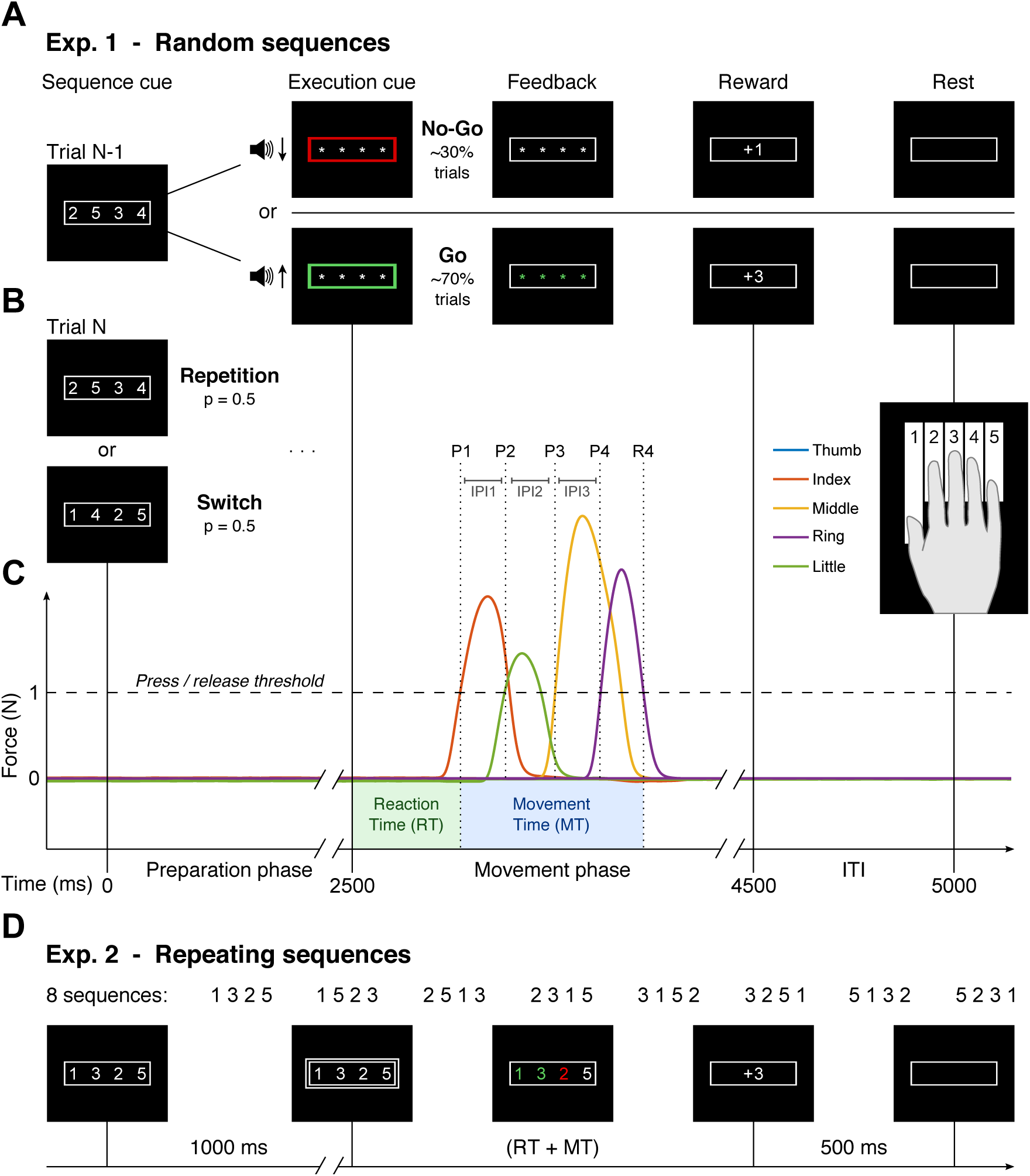
The discrete sequence production (DSP) task. **A.** Exp. 1 example trial: a sequence cue (white numbers on the computer screen) is followed by a production cue (outline changes color, numbers are masked). Online visual feedback about keypresses was given during the movement phase (green asterisks for correct presses, red for incorrect presses), followed by reward points depending on performance. 30% of the trials in a block were No-Go trials (red outline + low-pitch sound, top), 70% were Go trials (green outline + high-pitch sound, bottom). **B.** The next trial could be either a Repetition of the same sequence (0.5 probability), or a Switch to a new sequence. **C.** Example trial in Exp. 1 with the following trial timing: preparation phase: 2.5 sec; movement phase: 2 sec; ITI: 0.5 sec. Dashed horizontal line indicates force threshold (1 N) to determine the moment of each keypress and release (dotted lines). P1 = press of first key; R4 = release of fourth key; IPI1 = first inter-press interval. Total time (TT) = RT + MT. **D.** Exp. 2 design: 8 repeating sequences, trial structure, and timing. The go signal here is given via white box around the sequence cue.

Visual stimuli to instruct one sequence of finger presses were presented on a computer monitor and consisted of a string of 4 numeric characters displayed in white on black background (Sequence cue) and framed by a white rectangle (Fig. 1A; character height 1.5 cm, visual angle approx. 2°).

### Task

We used a discrete sequence production (DSP) task in which participants were required to produce sequences of keypresses with the five fingers of their right hand (Fig. 1A). Each sequence was cued by 4 numbers ranging from 1 to 5, instructing which fingers had to be pressed (e.g., 1 = thumb, 2 = index, … 5 = little). The sequence had to be produced by sequentially pressing the fingers corresponding to the numbers on the screen, from left to right. On each trial, participants were presented with a 4-item sequence and asked to prepare for the corresponding finger presses (preparation phase). After a fixed delay of 2.5 seconds, an audio-visual production cue would mark the beginning of the movement phase (a fixed 2 seconds). On Go trials, the production cue was a green frame accompanied by a high-pitch tone (Fig. 1A, bottom), indicating that participants had to perform the planned sequence of finger presses as quickly and accurately as possible (Go condition). On other trials, the production cue was a red frame accompanied by a low-pitch tone (Fig. 1A, top), instructing the participants to remain as still as possible without pressing any key until the end of the movement phase (No-Go condition). To encourage sequence pre-planning before the production cue, at the beginning of the movement phase the sequence cue was replaced by 4 asterisks masking the numbers. Moreover, the sequence of keypresses had to be completed within 2 seconds from the production cue (timeout error after that). With each keypress, the corresponding asterisk turned either green (correct press) or red (wrong press). Performance was evaluated in terms of both execution speed and press accuracy. Speed was defined in terms of total time (TT), which consisted of the reaction time (RT: from the onset of the Sequence cue to the first keypress) plus the movement time (MT: from the onset of the first keypress, P1, to the release of the last keypress, R4). A single press error invalidated the whole trial, so accuracy was calculated as percent error rate (ER) per block of trials (number of error trials / number of total trials x 100). At the end of the movement phase, during the 500 ms inter-trial interval (ITI), participants were presented with performance points appearing in place of the asterisks.

### Feedback

To motivate participants to improve in speed (TT = RT + MT) and accuracy (1 - ER) of sequence production, we gave participants performance feedback on each trial. The performance score was based on the following point system: −1 points for timing errors (i.e., anticipation of the production cue, or movement initiation in No-Go trials); 0 points for correct timing but wrong finger press (any one wrong keypress); +1 points for correct timing and press (i.e., movement initiation in Go trials, or no movement in No-Go trials); and +3 points for correct timing, correct press, and TT 2% or more faster than TT threshold. TT threshold would decrease by 2% from one block to the next if both of the following performance criteria were met: median TT in the current block faster than best median TT recorded hitherto, and mean ER in the last block < 25%. If either one of these criteria was not met, the thresholds for the next block remained unchanged. At the end of each block of trials, the median TT, mean ER, and points earned were displayed to the participants. At the end of the session, monetary compensation corresponded to the amount of performance points accumulated (points < 750 = 10 $; 750 ≤ points < 1000 = 12 $; points ≥ 1000 = 15 $).

Penalizing timing errors (−1 points) more than press errors (0 points) might have made participants more cautious and increased their RTs. Thus, to encourage full preparation of the sequence, for the last 20 participants of Exp. 1 we gave equal weight to timing and press errors (both 0 points). To check whether the penalty for timing errors affected RT performance, out of the 45 participants not excluded from data analysis, we compared the reaction times of participants who received the penalty (N = 27) and those who did not (N = 18). An independent samples *t*-test showed no statistical difference in RTs between the two groups (with or without penalty for timing errors), suggesting that participants adopted a similar strategy regardless of the penalty (penalty group: 443 ± 15 ms; no-penalty group: 453 ± 19 ms; difference: −10 ± 23 ms; *t*_43_ = −0.437, p = 0.664).

The scoring system in Exp. 2 was identical to the one in Exp. 1 without any additional penalty for eventual timing errors (0 points). Participants in Exp. 2 were paid a flat hourly rate (7 $), regardless of the specific amount of points accumulated.

### Design

#### Exp. 1

To investigate the nature of the repetition effect, we used a 2-by-2 design independently manipulating whether a particular sequence was repeated or whether the previous trial was a Go or No-Go trial. Sequences in Exp. 1 were randomly determined (see below). On any given trial, there was a 0.5 probability that the sequence was the same as the previous trial (Repetition) or that it was different (Switch; Fig. 1B). Independently, we varied whether each trial was a Go trial (70%) or a No-Go trial (30%). We designed a majority of the trials to be go-trials to encourage full sequence pre-planning before the production cue. The order of trials was randomly interleaved, creating all possible combinations of the factors repetition type and execution type of the previous trial. Note that, given that the trial structure was kept fixed across all experimental conditions (i.e., 2.5 sec preparation phase + 2 sec movement phase + 0.5 sec ITI), there was no difference in time elapsed after a Go, or No-Go trial. Each block was composed of 48 trials (12 repetitions for each of the 4 sequences), and participants underwent 1 session of 12 blocks each. In order to limit strong learning effects that might lead to ceiling performance, for each block of trials, we randomly selected four different 4-item sequences from a large pool of all permutations with repetition of the numbers 1 to 5, taken 4 items at a time. Moreover, to keep sequences of a similar level of difficulty, we removed from the permutation pool all sequences in which any number repeated (i.e., each number could only appear once per sequence), or that included “runs” (more than 2 fingers in either increasing or decreasing order; e.g., 1-2-3, or 3-2-1).

#### Exp. 2

To explore how sequence-specific learning affects sequence repetition, we designed a second experiment where one set of 8 sequences remained fixed over time. Participants underwent training for two consecutive days to ensure the development of enough sequence-specific learning. However, for the purposes of this study, we will not be examining consolidation effects, which will be discussed in future work. We used 8 4-item sequences including all fingers of the right hand except for the ring finger. The sequences were selected according to the following criteria: 1) each finger was used only once per sequence; 2) each finger started 2 of the 8 sequences; 3) each finger was pressed in every ordinal position twice across sequences; and 4) no more than 2 neighboring fingers pressed in a row (i.e., as in Exp. 1, we excluded “runs”).

In contrast to Exp. 1, Exp. 2 did not contain any No-Go trials and the preparation phase was shortened to a fixed 1 s. Also, the production cue was presented only visually (white box around the sequence cue), the sequence cue was not masked, and the duration of the movement phase was not fixed (i.e., TT dictated the actual duration of the trial, with the ITI occurring right after the last keypress). Finally, sequence repetition was not randomized, but counter-balanced across sequences. Each sequence was executed from a minimum of once (i.e., a Switch) to a maximum of five times in a row (i.e., executing once and repeating four times). To ensure a comparable number of trials per each repetition condition, we manipulated the proportion of same-sequence executions in a row as follows: 0.33 one-execution trials (Switch), 0.22 two-executions trials (One-repetition), 0.22 three-executions trials (Two-repetitions), 0.11 four-executions trials (Three-repetitions), 0.11 five-executions trials (Four-repetitions). Each of the 8 sequences was presented in each repetition condition the same number of times (balanced design across sequences), and the factors sequence type and repetition condition were then pseudo-randomized within a block of trials. In addition, unbeknownst to the participants, we included a variable number (from 1 to 4) of one-execution trials (Switch) as dummy trials at the beginning (assuming warm-up) and end (assuming tiredness) of each block, which were subsequently excluded from data analysis. Overall, this led to a final proportion of 0.59 repetition trials (across repetition conditions), and 0.41 switch trials. Each experimental block consisted of 50 trials (including dummy trials), and participants performed 12 blocks (∼ 4-5 min each) per experimental session per day (i.e., 24 blocks in total per participant). No explicit information about the sequence types, or instruction to memorize the sequences, was given. Nonetheless, the extensive repetition (more than 130 trials per sequence) ensured that participants would learn the finger transitions associated with each sequence, usually already by the end of the first testing day.

### Data analysis

Data were analyzed offline using custom code written in MATLAB (The MathWorks, Inc., Natick, MA). Statistical analyses for assessing movement repetition effects on reaction times (RT) and sequence movement time (MT) included two-tailed paired-samples t-tests (Repetition vs. Switch), and 2-by-2 within-subject repeated measures ANOVAs with factors repetition type (Repetition / Switch) and previous trial type (No-Go / Go). Error trials (both timing and press errors), No-Go trials, and dummy trials (Exp. 2 only, see Design section) were excluded from data analysis. For visualization purposes only, data were normalized by subtracting from each data point each participant’s mean and adding back the grand mean of the group. Statistical analyses, computed on raw data, were not affected by this normalization procedure.

## Results

### Sequence repetition reduces both reaction and movement times

Our experiment was designed to test whether there are short-term (i.e., trial-to-trial) benefits for the repetition of sequential movements. We compared trials in which the movement sequence was the same as on the previous trial (i.e., Repetition trials) to trials preceded by a different sequence (i.e., Switch trials). We found that RT improved upon repetition of the same sequence (Fig. 2A). Participants could react more quickly to the Go cue when then previous trial contained the same sequence (paired-samples t-test, *t*_44_ = 2.890, *p* = 0.006). Notably, this RT advantage was present even though participants had more than enough time (2.5 s) to finish pre-planning the 4-item sequence before the production cue (Ariani and Diedrichsen 2019). These results extend previous insights by Mawase et al. (2018) by showing that repetition facilitates the triggering of a planned movement, an effect that cannot simply be accounted by improved identification of the visual stimuli, or a bias in selection processes. Repetition also accelerated sequence production, as indicated by a significant repetition effect on sequence MT (Fig. 2C). On Repetition trials, the sequence was performed 36 ms (± 5 ms) faster than on Switch trials, a robust effect across participants (paired-samples t-test, *t*_44_ = 7.473, *p* = 2.330e-09).

**Figure 2.**
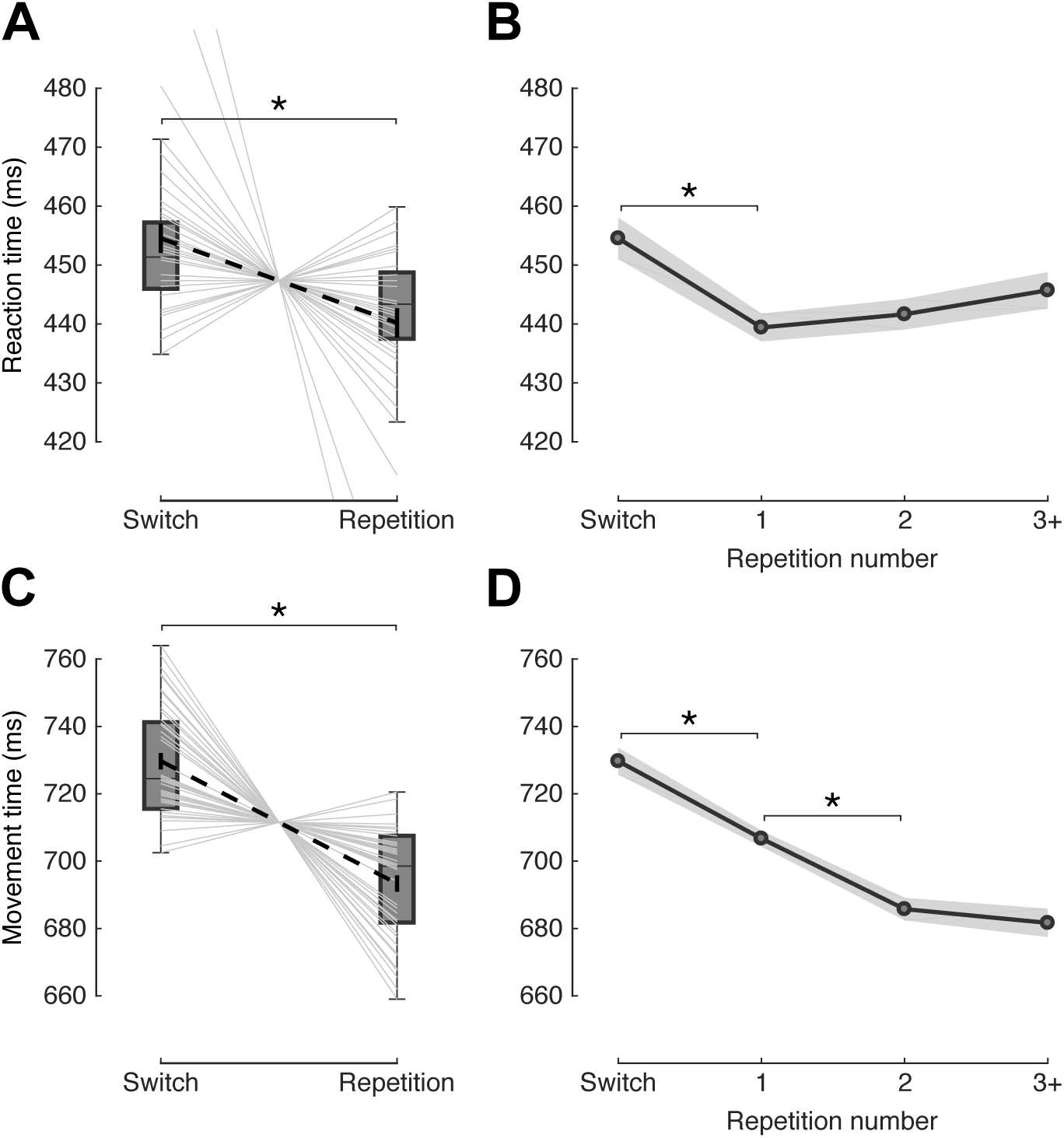
Immediate repetition leads to better performance. **A.** Distribution of median reaction times (RT) separately for Switch and Repetition trials. Light-gray lines represent individual participants. Dashed black line shows group mean across conditions, with relative standard error. **B.** Mean RT as a function of repetition number (1 means that a sequence was performed twice in a row, 3+ means the average of a sequence being repeated 4 or more times in a row). Shaded areas represent between-subject standard error of the mean. **C.** Distribution of median sequence movement times (MT) separately for Switch and Repetition trials. Other conventions as in panel A. **D.** Mean sequence MT as a function of repetition number. Other conventions as in panel B. *p < 0.05, two-tailed paired-samples t-test.

Next we asked whether the repetition effect would increase with further repetitions of the same sequence. For RT, the repetition effect was limited to the first repetition (Switch-Rep1 difference: 15 ± 4 ms; *t*_44_ = 3.753, *p* = 5.081e-04; Fig. 2B). After that, no further RT advantage was observed for successive sequence repetitions (Rep1-Rep2 difference: −2 ± 4 ms; *t*_44_ = −0.537, *p* = 0.594; Rep2-Rep3+ difference: −4 ± 3 ms; *t*_44_ = −1.138, *p* = 0.261). In contrast, for MT the improvements were not limited to the first repetition (Switch-Rep1 difference: 23 ± 4 ms): a second repetition (i.e., performing the same sequence three times in a row) was almost nearly as beneficial to further reduce sequence MT (Rep1-Rep2 difference: 21 ± 4 ms; *t*_44_ = 4.712, *p* = 2.478e-05; Fig. 2D). After the second repetition, performance appeared to reach a plateau (Rep2-Rep3+ difference: 4 ± 6 ms; *t*_44_ = 0.705, *p* = 0.484).

Importantly, faster RT and MT in Repetition trials did not come at the cost of decreased accuracy. In fact, the opposite was true: accuracy increased from 81.8 ± 1.1% correct trials in Switch trials to 85.6 ± 1.2 % in Repetition trials (paired-samples t-test *t*_44_ = −5.532, *p* = 1.637e-06). Moreover, timing errors (i.e., false starts by anticipation of the production cue) decreased from 4 % to 2.9 % (paired-samples t-test *t*_44_ = 2.777, *p* = 0.008).

Overall, our results suggest that repetition of a sequence improves both the initiation of a pre-planned movement, as well as the speed by which the repeated sequence can be performed. The accuracy advantage proved that this effect did not arise at the expense of reduced execution accuracy.

### The repetition benefit arises from improved online planning

The results so far indicate that sequence repetition improves initiation (RT) and movement (MT). Should this be taken as an indication that repetitions improve execution-related, rather than planning-related processes? Not necessarily so. In a previous study we have demonstrated that sequence MT (the time from first to last keypress) is not only a function of motoric processes, but is also strongly influenced by the speed of online planning (Ariani and Diedrichsen 2019). Even for short sequences, only the first 2-3 keypresses can be fully pre-planned, whereas later movements appear to be planned online, that is during of the execution of the beginning of the sequence. If movement repetition facilitates online planning, this effect should therefore be more prevalent in latter parts of the sequence. If, however, movement repetition facilitates execution processes, it should influence the speed of all presses in the sequence, no matter if these are performed in the beginning or later.

To examine this issue, we inspected the 3 inter-press intervals (IPIs) between the onsets of the 4 keypresses separately. The second transition was the slowest, while the first and last transition were nearly equally fast (Fig. 3A). This indicates a “2-and-2” rhythm, in which each 4-item sequence begins with two quick presses, followed by a brief pause, and then again by two quick presses. Given that the sequences changed randomly from block to block, all possible finger transitions could occur with equal probability at each position of the sequence. Therefore, this effect cannot be explained by biomechanical factors (e.g., some transitions being harder than others). Instead, the pattern or results suggests a clear influence of online planning: the first two keypresses can be fully pre-planned and can therefore be executed quickly; then execution needs to slow down until online planning of the remaining two keypresses is finished.

**Figure 3.**
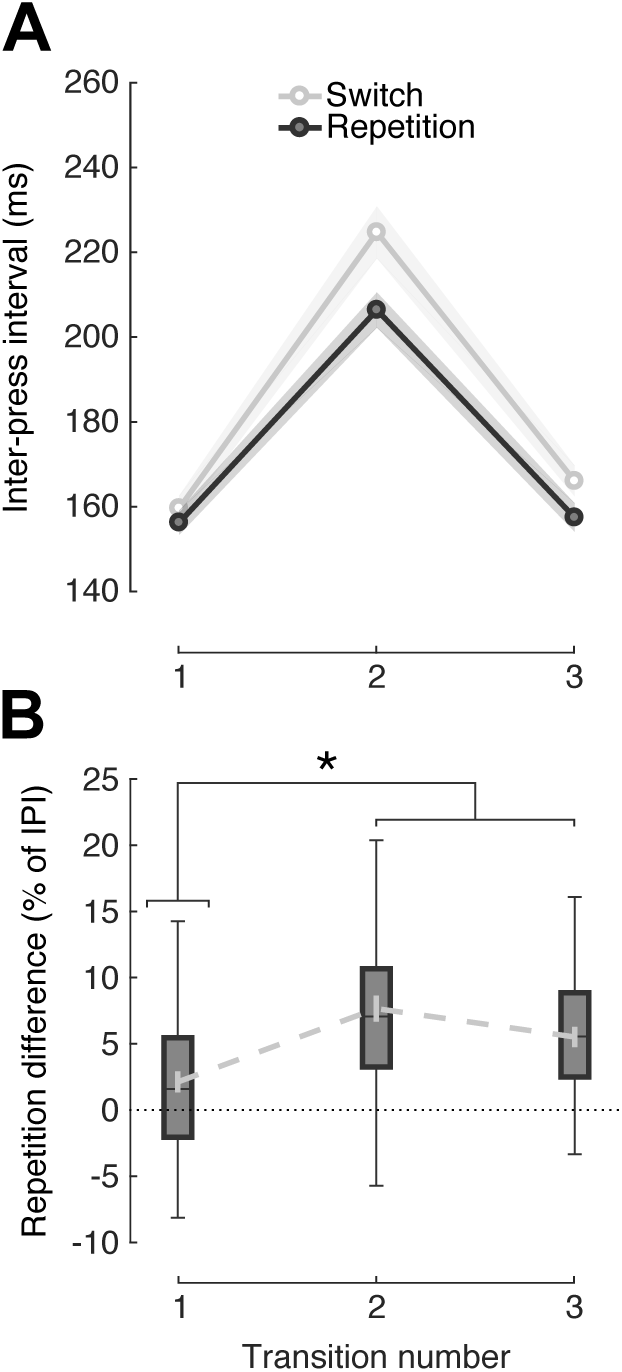
Sequence repetition benefits online planning of repeating sequence elements. **A.** Mean inter-press intervals (IPI) as a function of transition number, separately for Switch (light) and Repetition (dark) trials. Shaded areas represent between-subject standard error of the mean. **B.** Same data as in A but normalized by mean IPI for each transition separately Dashed gray line denotes group mean across conditions, with relative standard error. *p < 0.05, two-tailed paired-samples t-test.

Importantly, we found that the repetition effect was most pronounced on the second transition (Switch-Repetition difference, 2^nd^ vs 3^rd^ transition: 10 ± 3 ms; *t*_44_ = 3.205, *p* = 0.003; Fig. 3A). After normalizing the Switch-Repetition difference by mean IPI of each transition, we found that the repetition benefit, expressed as percentage of average IPI (Fig. 3B), was significantly greater on the second and third transitions than on the first transition (*t*_44_ = 5.380, *p* = 2.729e-06). Taken together, the pattern of IPIs is consistent with the view that repetition affects sequence movement times by accelerating processes related to online planning and does not speed up the actual production of individual keypresses.

### Repetition effects may require actual movement experience

So far, the results suggest that sequence repetition accelerates subsequent pre- and online planning processes. Next, we addressed the question of whether pre-planning of a sequence would be sufficient to produce a benefit on subsequent trials RT, or whether the execution of the sequence (involving initiation, motor processes, and online planning) may be required. For this purpose, we compared Switch and Repetition trials separately for whether the previous trial (N-1) had been a Go, or a No-Go trial. Our logic was that if the repetition effect resulted from stimulus processing, selection, and pre-planning, we should see a repetition benefit even if the previous trial had been a No-Go trial. Conversely, if the repetition effect requires the initiation or execution of the sequence, then we should only observe it when the previous trial had been executed (i.e., N-1 was a Go trial). To avoid repetition trials which were preceded both by a Go and by a No-Go trial of the same sequence, we restricted this analysis to the first repetition of a sequence (i.e., max two executions in a row).

We found that the repetition effect on RT was significant when the previous trial had been a Go (*t*_44_ = 4.534, *p* = 4.421e-05), but not when it had been a No-Go (*t*_44_ = 0.986, *p* = 0.330; Fig. 4A). Importantly, the interaction between repetition type (Switch vs. Repetition) and previous trial type (No-Go vs. Go) was significant (2-by-2 within-subject ANOVA, *F*_1,44_ = 5.303, *p* = 0.026). The same pattern of results was observed for MT (Fig. 4B): significant repetition effect only on N-1 Go trials (*t*_44_ = 5.464, *p* = 2.055e-06), and significant interaction between repetition and previous trial type (*F*_1,44_ = 4.898, *p* = 0.032). As expected, the effect was also visible when we split up the MT into IPIs (Fig. 4C). Only for the second transition did we find a significant interaction between repetition type and previous trial type (*F*_1,44_ = 4.271, *p* = 0.044).

**Figure 4.**
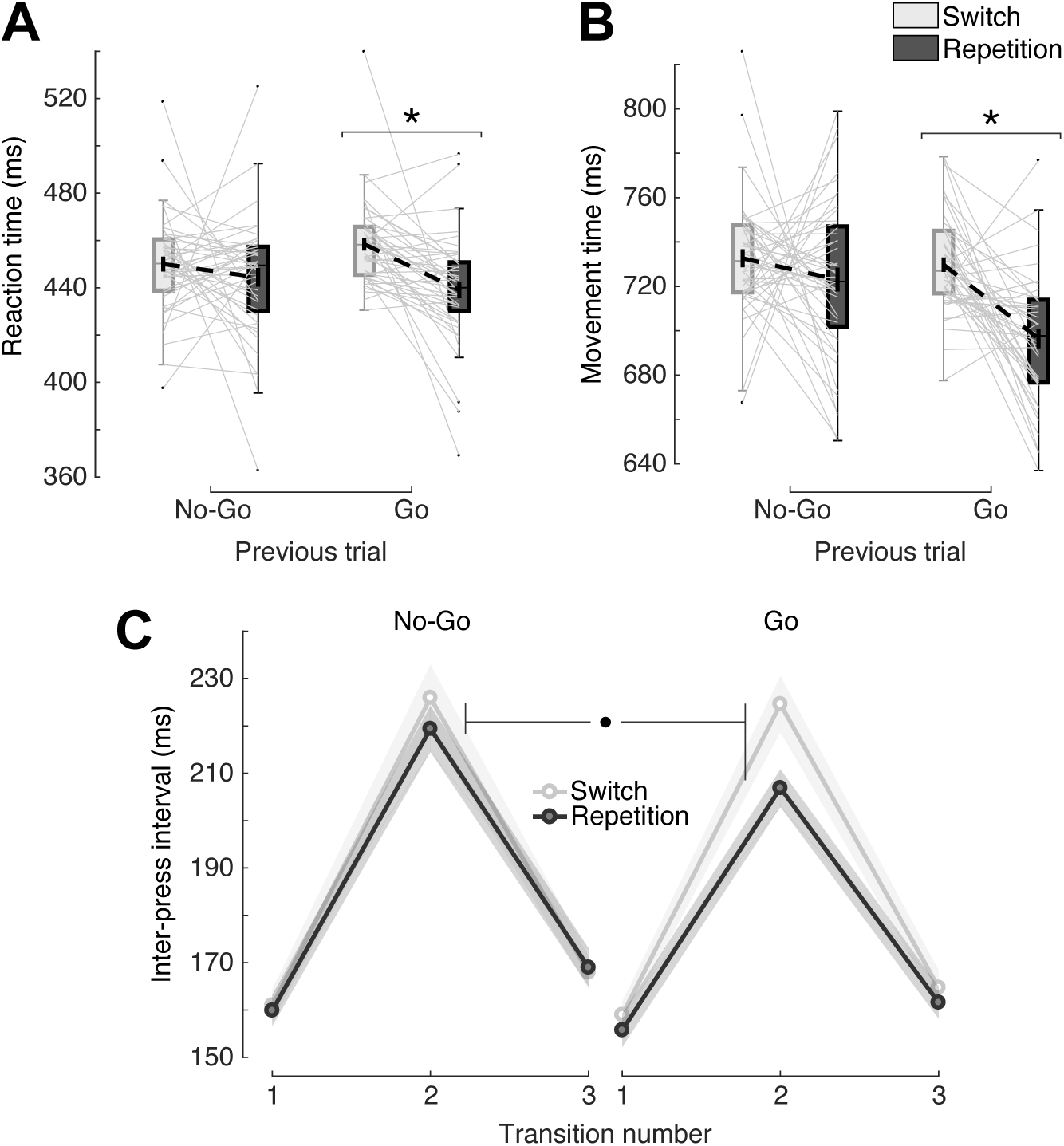
The repetition effect appears to require movement experience. **A.** Distribution of median movement times (MT) separately for Switch (light) and Repetition (dark) trials as a function of whether the previous (N-1) trial was a Go, or a No-Go trial. To avoid contamination between Go and No-Go trials in long repetition chains, selected trials were restricted to a maximum of one repetition. Solid light-gray lines represent individual participants. Dashed black lines represent group means across conditions. Black dots are considered group outliers. **B.** Distribution of median reaction times (RT) separately for Switch (light) and Repetition (dark) trials as a function of whether the previous (N-1) trial was a Go, or a No-Go trial. Other conventions as in panel A; **C.** Mean inter-press intervals (IPI) as a function of transition number, separately for Switch (light) and Repetition (dark) trials and split by whether the preceding trial was a No-Go (left) or a Go (right) trial. Shaded areas represent between-subject standard error of the mean. *p < 0.05, two-tailed paired-samples t-test; ^**•**^p < 0.05, for interaction in 2-by-2 within-subject repeated measures ANOVA.

Again, faster reaction and movement in repetition trials did not trade-off with sequence execution accuracy. The strong and consistent improvements in accuracy after a repetition (*F*_1,44_ = 26.698, *p* = 5.543e-06) was not different depending on whether the previous trial had been a Go or a No-Go trial (*F*_1,44_ = 0.159, *p* = 0.691), nor it was the decrease in timing errors (main effect of repetition, *F*_1,44_ = 6.238, *p* = 0.016; interaction between repetition and previous trial execution, *F*_1,44_ = 0.190, *p* = 0.665).

Taken together, our results are consistent with the view that the repetition effect relies on the experience of performing, or at least initiating the execution of a sequence. The act of processing the visual stimuli, selecting, and pre-planning the upcoming sequence movements (all processes that we assumed would be performed on No-Go trials as well) was not sufficient to obtain faster RT or MT on Repetition trials. Therefore, the benefit of rehearsal in a motor sequence task appears to require processes that are only activated when the movement is actually initiated or executed.

### Repetition improves the speed of even the fastest movements and participants

Immediate movement repetition makes participants faster at initiating and producing a motor sequence. This effect may have been caused by an improvement in movement speed across the board – that is, even the fastest trials should get even faster after repetition. Alternatively, the effect could have been caused by the fact that repetition makes slow trials less likely: such slow trials may be the result of errors in planning, lack of concentration or focus on the task. In other words, repetition may simply ease the computational burden on the subject and make suboptimal trials less likely, without actually affecting the top speeds in sequence production. To investigate this idea, we divided the whole distribution of sequence movement times into 11 bins for each repetition condition and participant separately. This analysis was performed separately for trials for which the previous trial (N-1) was a No-Go or a Go trial (Fig. 5A). For Go trials, the repetition effect was present for the whole range of sequence execution speeds. Importantly, this was true even for the fastest MTs: the mean repetition difference for the fastest bin was 41 ± 19 ms (one-sample *t*-test vs. zero *t*_44_ = 2.124, *p* = 0.039).

**Figure 5.**
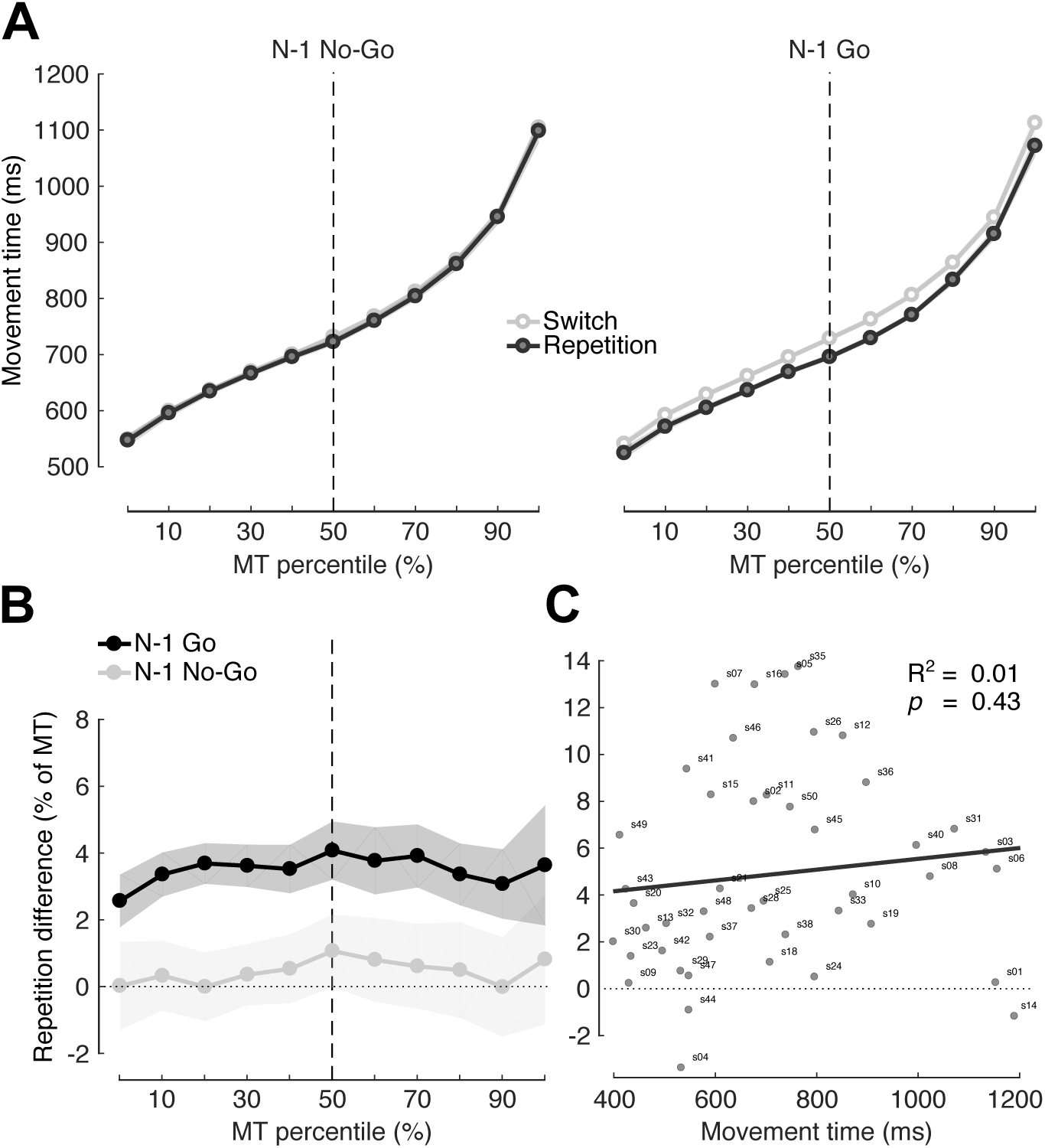
The repetition benefit remains constant across a wide range of movement speeds. **A.** Sequence MT as a function of MT percentile (11 bins), separately for Switch (light) and Repetition (dark) trials, and by previous trial No-Go (left), or Go (right). Dashed vertical line denotes the MT bin that includes the median MT. Shaded areas represent between-subject standard error of the mean. **B.** Repetition difference normalized by median MT for each bin separately as a function of MT percentile, and by previous trial No-Go (light), or Go (dark). Other figure conventions are the same as in A. **C.** Repetition difference normalized by median MT plotted against MT for each participant. Solid line represents linear regression. Dotted line as landmark for lack of repetition difference. R^2^ = proportion of variance in repetition difference explained by execution speed.

To better quantify this difference, we plotted the repetition difference normalized by the MT of the corresponding percentile bin (i.e., [Switch MT – Repetition MT] / Overall MT for each bin; Fig. 5B). Again, this analysis confirmed that for N-1 Go trials the percentage repetition effect was significant (one-sample *t*-test vs. zero across all percentiles, *t*_44_ = 4.792, *p* = 1.908e-05) and did not differ across the range of speeds (paired-samples *t*-test first vs. last percentile, *t*_44_ = −0.516, *p* = 0.609). As expected, for N-1 No-Go trials the effect was not significant (*t*_44_ = 1.064, *p* = 0.293). Thus, the repetition benefit was not simply a consequence of preventing occasionally slow executions (i.e., an attentional effect). Rather, after a repetition, participants were more likely to beat their currently best speed.

Lastly, we investigated whether repetition was equally beneficial for all subjects, or only for participants that were relatively slow performers to begin with. Indeed, for participants that were already faster overall, the benefit of repetition may not have constituted a large proportion of their movement time. For each participant, we plotted the median Switch-Repetition difference normalized by MT as a function of median sequence movement time (Fig. 5C). The correlation between sequence production speed and size of the repetition effect was not significantly different from zero (r = 0.119, *t*_44_ = 0.790, *p* = 0.434), so we found no evidence that inter-subject variability had a strong influence on the relative repetition benefit. Overall, we showed that movement repetition enables faster sequence MT across the board – both for fast and slow trial, and for fast and slow participants.

### Does sequence-specific learning affect the repetition benefit?

In Exp. 1, sequences varied randomly from block to block, effectively preventing participants from learning a specific set of keypress transitions. Therefore, it is likely that overall performance improvements largely reflected sequence-general learning processes (e.g., faster single-item selection, or task familiarization). But how does sequence-specific learning (i.e., extensively practicing a fixed set of sequences) change the repetition effect? One possibility is that practice may diminish, and eventually erase, the repetition effect. This would be consistent with our previous finding that sequence-specific learning reduces the time that participants need to pre-plan and online-plan the sequences (Ariani and Diedrichsen 2019).

To test this hypothesis, we designed a second study (Exp. 2) in which a set of 8 sequences was kept constant throughout the experiment (12 blocks of 50 trials each). We still randomly varied, however, whether a sequence would repeat or change between trials. We then analyzed how the repetition effect changed over the course of learning. For illustrative purposes, we re-analyzed the results of the first experiment (Exp. 1, N = 45, sequence-general; Fig. 6A-6B, left column) in the same format as the results for the second experiment (Exp. 2, N = 39, sequence-specific; Fig. 6A-6B, right column) and focused on the comparison between the first day of both experiments.

**Figure 6.**
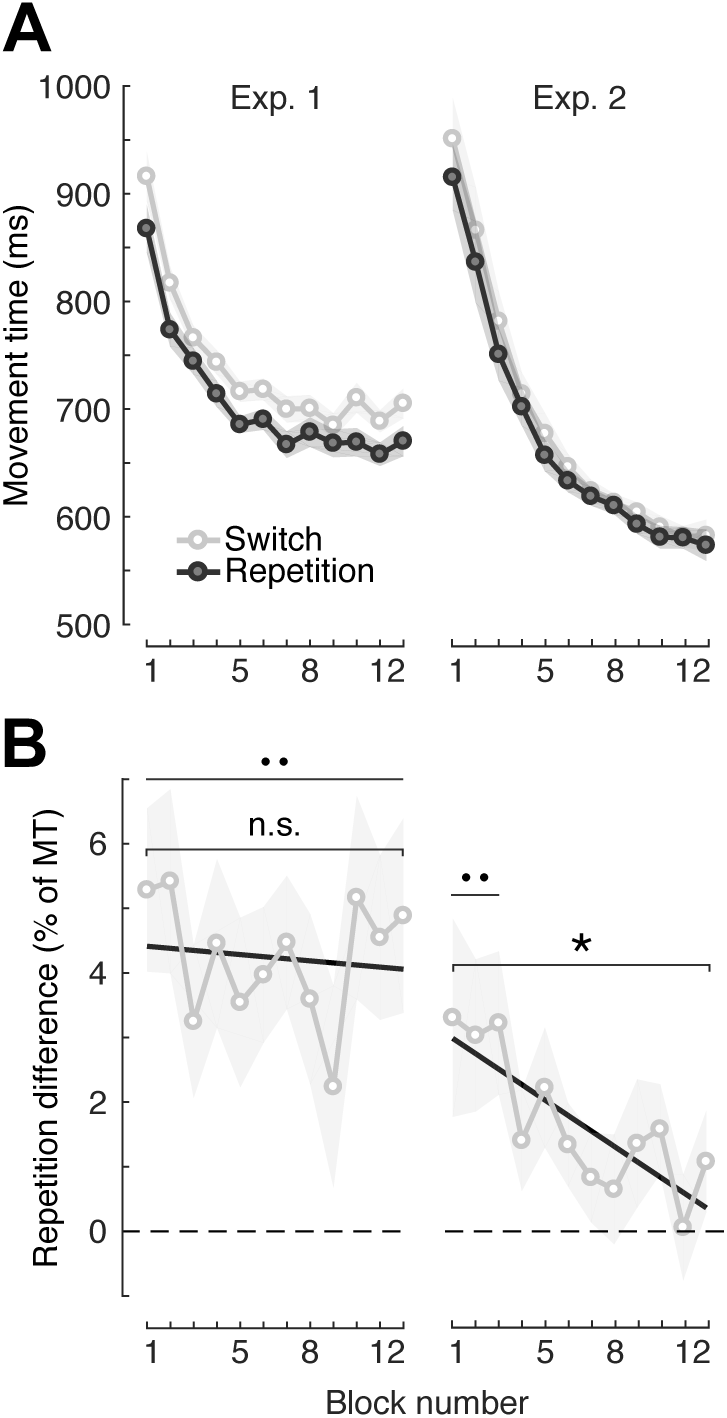
Sequence-specific learning reduces the repetition effect. **A.** Sequence movement time separately for Switch (light) and Repetition (dark) trials as a function of block number in Exp. 1 (left) and Exp. 2 (right). Shaded areas represent between-subject standard error of the mean. **B.** Switch-Repetition difference normalized as percentage of MT for each block in Exp. 1 (left) and Exp. 2 (right). Shaded areas represent between-subject standard error of the mean. Black solid line represents linear regression line. Dashed horizontal line indicates absence of repetition effect. *p < 0.05, two-tailed paired-samples t-test; ^**• •**^ p < 0.05, two-tailed one-sample t-test vs zero difference.

In Exp. 1, where sequence-specific learning was nearly impossible, a clear repetition benefit was present from the first block (Fig. 6A, left; Switch-Repetition difference: 49 ± 12 ms, *t*_44_ = 3.973, *p* = 2.600e-04) and remained roughly constant despite sequence-general learning, until the last block (last block Switch-Repetition difference: 35 ± 11 ms, *t*_44_ = 3.315, *p* = 0.002; Fig. 6A, left). Importantly, once adjusted by overall MT for each block (Fig. 6B, left), there was no difference in repetition benefit from block 1 to block 12 (paired-samples *t*-test, *t*_44_ = 0.214, *p* = 0.832).

In Exp. 2, we found that the repetition benefit was present in the first few blocks (one-sample t-test between mean blocks 1-3 vs. zero difference, *t*_38_ = 2.424, *p* = 0.020; Fig. 6B, right) despite the overall improvement in sequence execution speed (Fig. 6A, right). However, this effect quickly vanished after a few blocks of practice (repetition difference on block 4, *t*_38_ = 1.806, *p* = 0.078; Fig. 6B, right).

These results show that the repetition effect gradually decreases with practice. Importantly, sequence-specific, but not sequence-general learning was associated with such decrease.

## Discussion

In two behavioral experiments, we establish that repeating a sequence of movements led to immediate improvements in reaction times and movement speed (Fig. 2) without any associated cost in performance accuracy. The repetition benefit during sequence production was largest in the middle part of a sequence (Fig. 3), and was absent for the first two presses, suggesting that repetition did not affect execution-related processes (which should be involved in all presses), but rather online planning (which was most relevant in the middle of the sequence). The finding that sequence repetition facilitated online planning was consistent with the observation that the repetition effect decreased with sequence-specific, but not sequence-general practice (Fig. 6). That is, once sequence-specific learning reduced the role of online planning, the benefit of repetition disappeared. Finally, we observed that repetition-related improvements only occurred for the trials that had been preceded by sequence production (which involves movement initiation, execution, and online planning), suggesting that action selection and pre-planning may not be sufficient to drive the repetition effect (Fig. 4-5).

### Sequence-level repetition effects in motor sequence production

Repetition of a sequence of finger movements resulted in immediate improvements in speed and accuracy of sequence production. This indicates that repetition also affects processes governing the planning or execution of the entire movement sequence, rather than just individual movements. To understand the implications of this finding, it may be useful to consider our findings in the framework of neuronal state-spaces (Churchland et al., 2010, Fig. 7). In this framework, neural activity in movement-related brain regions (e.g., primary and pre-motor cortex) can be decomposed into neuronal dimensions representing the current movement (execution state-space) and neuronal dimensions representing the next upcoming movements (planning state-space). Pre-planning would be equivalent to bringing the neuronal population state into a specific location of planning state-space (Churchland et al. 2006b). Upon movement initiation, the neuronal state changes dramatically (Elsayed et al. 2016; Kaufman et al. 2016) and subsequently evolves mainly in the dimensions that span the execution state-space, generating the patterns required for producing muscular output. While neurons in the dorsal pre-motor cortex (PMd) likely contribute more to the planning state-space, neurons in the primary motor cortex (M1) contribute more to the execution state-space. However, although execution- and planning-related signals are mixed in these two regions, with many neurons responding to both processes (Alexander and Crutcher 1990; Prut and Fetz 1999; Riehle and Requin 1989), planning and execution processes can be kept from interfering with each other by using orthogonal neural dimensions of the same overlapping population of neurons (Kaufman et al. 2014).

**Figure 7.**
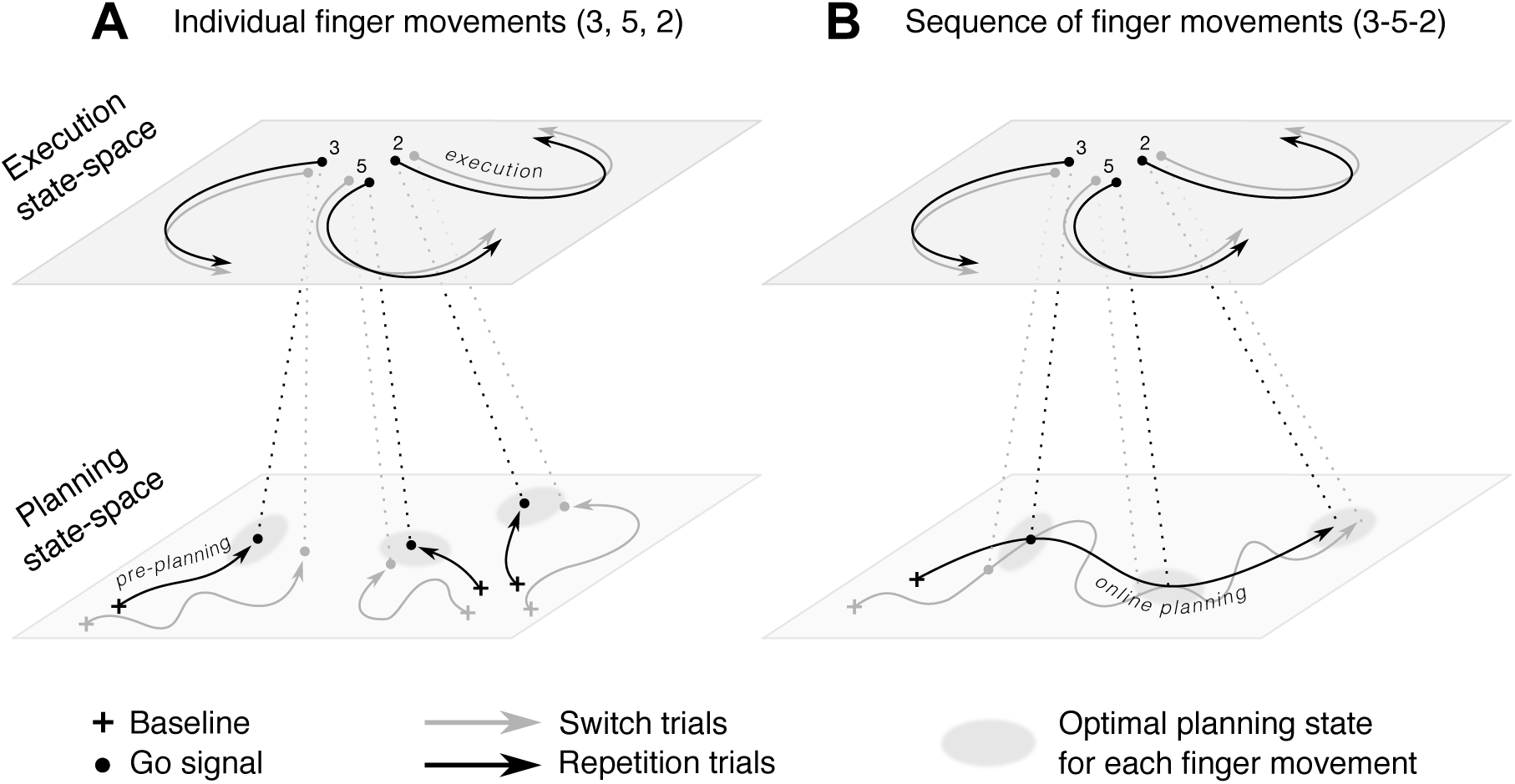
Conceptual visualization of planning and execution in a neural state-space framework. Changes in the pattern of neuronal firing can be characterized by movements of the neuronal state in a low-dimensional state-space. During the preparation phase, these changes mostly occur in the planning state-space (lower plane), whereas during the movement phase, these changes are more pronounced in the execution state-space (upper plane). Different finger movements (numbers, see Fig. 1) are characterized by a unique pattern in the planning state-space, and a unique trajectory in the execution state-space. Dotted lines indicate temporal correspondence between state-space events across planning and execution planes. **A.** Individual finger movements. During pre-planning (after stimulus onset), the neural state trajectory moves from a baseline location (cross) towards the optimal planning state (gray spotlight) until the go signal (dot) triggers execution dynamics. On repetition trials the correct planning state would be reached more quickly and with higher accuracy, enabling faster movement initiation when the go signal is given. **B.** Sequence of finger movements. Note that in this scenario what improves upon repetition is the neural state trajectory on the planning plane (i.e., online planning), leaving neural dynamics unchanged on the execution plane.

In this framework, repetition effects for individual finger movements would be caused by the fact that the correct pre-planning state can be reached faster and with more accuracy after repetition (Mawase et al. 2018), possibly by lingering activity in the planning state-space (Fig. 7A). Now consider the production of a short sequence. Here, the planning-related neural state needs to traverse multiple locations, each triggering the corresponding elementary movements in the execution state-space (Fig 7B). During movement, the neural state in the planning state-space would already start to plan the next movement (i.e., online planning). If movement repetition simply primed one location in the planning state-space, then any advantage of pre-planning the first individual movement element would be washed out after completing the sequence. Thus, the presence of a repetition effect at the level of sequences indicates that movement repetition primes the entire pathway through the planning state-space. This is consistent with the hypothesis of an intermediate level between movement selection and execution (Diedrichsen and Kornysheva 2015).

### RT advantage does not reflect improved stimulus processing or action selection

In a previous study we found that benefits of a prolonged preparation phase asymptote after ∼1.5 seconds (Ariani and Diedrichsen 2019). Thus, by using a delayed-response paradigm, we can be relatively confident that our RT measure mainly reflects the initiation of a pre-planned response, as processes of stimulus identification and action selection should have been completed during the preparatory delay (2.5 s). Thus, our study provides stronger evidence than Mawase et al. (2018), who used a Free-RT/Timed-response paradigm, that the repetition benefit on RT cannot be explained by faster perceptual processing of the target stimuli. By masking the sequence cue at the moment of the go signal, we purposely encouraged participants to complete the perceptual processing and response selection in the preparatory period. The faster response initiation after a repetition may indicate that the planning state was closer to the ideal state, which allowed for faster triggering of the desired sequence (Ames et al. 2014; Churchland et al. 2006a, 2006b; Michaels et al. 2018).

### Faster sequence production is due to more efficient online planning

Repetition accelerated not only RT, but also MT, for repeated sequences. Critically, a detailed analysis of the inter-press-intervals revealed that the transition between the first two keypresses, which was likely fully pre-planned, was not influenced by the repetition. Rather, the repetition advantage was observed on the second and, to some degree, third transitions. This finding is consistent with the view that repetition benefits arose as a consequence of facilitated online planning (Fig. 7B). Our current design cannot disambiguate whether this result was a consequence of participants splitting the 4-item sequences into 2 chunks, with online planning between the chunks, or whether the slowing down was caused by the necessity for continuous online planning in the middle of the sequence. Either way, sequence repetition shortens MT, not by accelerating how quickly individual movements can be executed, but by improving the speed in which sequence elements, be it chunks or individual presses, can be planned online.

### Sequence-specific learning gradually reduces the repetition effect

After one day of practice on a fixed set of sequences we observed behavioral improvements consistent with sequence-specific learning (Ariani and Diedrichsen 2019; Wiestler et al. 2014). This speed advantage went hand in hand with a decrease, and eventual disappearance, of the repetition effect. This result corroborates the interpretation that repetition benefits on sequence production come from improvements in online planning. More efficient online planning for known sequences allows for faster movement speeds, up to the point where participants are limited not by their ability to quickly plan the next response, but by the ability to motorically implement the response (Ariani and Diedrichsen 2019). When online planning ceases to be the main limiting factor, the repetition benefit disappears. An alternative and non-mutually exclusive interpretation is that sequences are planned and executed in movement chunks – in the case of our short 4-item sequences, 2 chunks of 2 keypresses each. After extensive training, participants could gradually learn to associate each sequence with a larger chunk of 4. This would enable them to quickly pre-plan the entire short sequence at once, and then execute it as one chunk, again removing the benefit of online planning during sequence repetition.

### Is pre-planning sufficient or is movement required to drive the repetition effect?

The repetition effect on MT was only present when the sequence was actually initiated and executed on the previous trial. Given our claim that the repetition is due to online planning, this finding would be expected, as sequence pre-planning alone would not move the neural state through the entire trajectory in the planning state-space (Fig. 7B). According to this view, neither the pre-planning of the initial part of the sequence, nor the execution of the individual sequence elements is enough to facilitate sequence production with repetition. Instead, it is revisiting the trajectory in the planning state-space that improves subsequent MT.

More surprisingly, pre-planning alone did not produce a repetition effect on RT. A priori, it was not obvious why executing a sequence would be required to observe a faster RT. In fact, if repetition facilitates response pre-planning (Mawase et al. 2018), one may have expected the persistence of the effect on RT. A potential explanation for this finding could be that, despite having enough time and information, participants did not fully pre-plan the response during the preparatory period. Indeed, reaction times were relatively long (∼400 ms) for triggering a pre-planned sequence. Perhaps participants used the time after the go signal to complete pre-planning. In this view, the completion of pre-planning up to and including movement initiation would be essential for the subsequent repetition benefit. Nonetheless, our experiment was designed to motivate participants to pre-plan the sequence well in advance during the delay. We masked the sequence cue so that they could not rely on it after the go signal. Go and No-Go trials were pseudo-randomly ordered, and we included a higher proportion of Go trials (70%), such that, more often than not, participants would be required to act on the pre-planned sequence. Finally, we rewarded participants on the sum of RT and MT, meaning that an easy way to earn more money would be to shorten RTs. Thus, it is hard to see how more complete pre-planning could be achieved. Instead, the act of initiating the sequence or online planning of the remainder of the sequence appear to be necessary to achieve faster RT on the next trial. However, how the processes related to pre-planning, initiation, and online planning interact with each other remains an open question.

### Conclusions

Our results show clear repetition effects for sequential movements, thereby extending previous findings that repetition speeds us the preparation of individual movements. The pattern of results is consistent with repetition facilitating trajectories through the preparatory neural state-space. While sequence production recruits widespread cortical sensorimotor areas (Kornysheva and Diedrichsen 2014; Wiestler and Diedrichsen 2013), our results would predict that the neuronal origin of repetition effects should not be found in the primary motor cortex, which mainly appears to be involved in the execution of individual movements (Yokoi et al. 2018; Yokoi and Diedrichsen 2019). Instead, we would expect to observe the effects of repetition in regions involved in (online) motor planning, such as dorsal premotor or superior parietal cortex.

## Acknowledgements

This work was supported by a James S. McDonnell Foundation Scholar award, an NSERC Discovery Grant (RGPIN-2016-04890), and the Canada First Research Excellence Fund (BrainsCAN). The authors wish to thank Jonathan A. Michaels for comments on the manuscript.

